# *qKW9* encodes a pentatricopeptide repeat protein affecting photosynthesis and grain filling in maize

**DOI:** 10.1101/847145

**Authors:** Juan Huang, Gang Lu, Lei Liu, Mohammad Sharif Raihan, Jieting Xu, Liumei Jian, Lingxiao Zhao, Thu M. Tran, Qinghua Zhang, Jie Liu, Wenqiang Li, Cunxu Wei, David M. Braun, Qing Li, Alisdair R. Fernie, David Jackson, Jianbing Yan

**Affiliations:** National Key Laboratory of Crop Genetic Improvement, Huazhong Agricultural University, Wuhan 430070, China; Cold Spring Harbor Laboratory, Cold Spring Harbor, NY, USA; Wimi Biotechnology Co., Ltd, 4th Floor, Kejizhuanhua building, No. 3 Meishan Road, Xinbei District, Changzhou City, Jiangsu Province, China; Jiangsu Key Laboratory of Crop Genetics and Physiology, Co-Innovation Center for Modern Production Technology of Grain Crops, Yangzhou University, Yangzhou 225009, China; Jiangsu Xuzhou Sweetpotato Research Center, Xuzhou, Jiangsu, China; Division of Biological Sciences, Interdisciplinary Plant Group, Missouri Maize Center, University of Missouri, Columbia, MO 65211, USA; Department of Molecular Physiology, Max-Planck-Institute of Molecular Plant Physiology, Am Mühlenberg 1, 14476 Potsdam-Golm, Germany

**Keywords:** Kernel weight, maize yield, QTL, photosynthesis, Cyclic electron transport, C-to-U editing, NDH complex

## Abstract

Kernel weight is an important yield component in maize that was selected during domestication. Many kernel weight genes have been identified through mutant analysis, and are mostly involved in the biogenesis and functional maintenance of organelles or other fundamental cellular activities. However, only a limited number of loci underlying quantitative variation in kernel weight have been cloned. Here we characterize a maize kernel weight QTL, *qKW9* and find that it encodes a DYW motif pentatricopeptide repeat protein involved in C-to-U editing of NdhB, a subunit of the chloroplast NADH dehydrogenase-like complex. In a null *qKW9* background, C-to-U editing of NdhB was abolished, and photosynthesis was reduced, suggesting that *qKW9* regulates kernel weight by controling the maternal source of photosynthate for grain filling. Characterization of *qKW9* highlights the importance of optimizing photosynthesis on maize grain yield production.

## INTRODUCTION

Maize (*Zea mays*) is one of the most important crops in the world, producing grain vital for our survival. Along with population growth, environmental deterioration, the decline of arable land and climate change challenge us to increase maize grain production. Therefore, the improvement of maize yield is of great importance to the sustainable development of human society.

The grain yield of maize is comprised of several components, including ear number per plant, kernel number per cob, and kernel weight. As an essential yield component, kernel weight is positively correlated with yield, and is determined during development by kernel size and the degree of kernel filling (Scanlon and Takacs, 2009). To dissect the genetic architecture of maize kernel weight, numerous studies have identified hundreds of quantitative trait loci (QTL) for kernel traits (www.maizegdb.org/qtl). However, only a few kernel size QTL have been cloned and studied, and some maize kernel weight genes have been identified as homologs of rice genes(Li et al., 2010a; Li et al., 2010b; Liu et al., 2015). In one large-scale QTL study in maize, 729 QTL regulating kernel weight-related traits were identified, and 24 of 30 genes were in, or tightly linked to, 18 rice grain size genes, suggesting conserved genetic architecture of kernel weight(Liu et al., 2017b). One example is *teosinte glume architecture1* (*tga1*), the causal gene underlying the change from encased kernels in the wild progenitor teosinte to naked kernels in maize(Wang et al., 2005; Wang et al., 2015). Reducing expression of *tga1* in maize by RNAi greatly increases kernel size and weight, suggesting that the removal of glumes from teosinte could release growth constraints, and provide more space to allow larger kernels to develop (Wang et al., 2015). Another kernel size gene, *ZmSWEET4c*, affects kernel weight in a different manner, with its product mediating sugar transport across the basal endosperm transfer cell layer, and shows signals of selection during domestication (Sosso et al., 2015). Recently a further gene, *BARELY ANY MERISTEM1d* (*ZmBAM1d*) was identified as an additional QTL responsible for kernel weight variation in maize (Yang et al., 2019).

Despite limited progress on our understanding of the quantitative variation in maize kernel weight, numerous kernel mutants have been identified (Neuffer and Sheridan, 1980; Clark and Sheridan, 1991). These mutants have been grouped into three categories: (i) defective kernel (*dek*) mutations, including empty pericarp (*emp*) mutants that affect both endosperm and embryo; (ii) embryo-specific (*emb*) mutations with healthy endosperm formation; and (iii) endosperm-specific mutations (McCarty, 2017). Mutants in categories i and ii have detrimental effects, leading to significant loss of kernel weight. Several of these maize kernel development genes have been identified. For instance, *EMP10*(Cai et al., 2017), *EMP11* (Ren et al., 2017), *EMP12* (Sun et al., 2019), *EMP16* (Xiu et al., 2016), *DEK35* (Chen et al., 2017), and *DEK37* (Dai et al., 2018) are involved in intron splicing of mitochondrial genes. In contrast, *MPPR6* functions in maturation and translation initiation of mitochondrial ribosomal protein subunit mRNA(Manavski et al., 2012). Mutations of these genes impair mitochondral function, leading to defective kernels. Other genes, including *EMP7* (Sun et al., 2015), *DEK10* (Qi et al., 2017), *DEK39* (Li et al., 2018), *PPR2263/MITOCHONDRIAL EDITING FACTOR29* (Sosso et al., 2012), and *SMALL KERNEL1* (Li et al., 2014) function in C-to-U editing of transcripts in mitochondria and chloroplasts.

Many kernel size genes encode pentatricopeptide repeat (PPR) proteins, a large family of RNA-binding proteins in land plants, with more than 400 members in *Arabidopsis* (*Arabidopsis thaliana*), rice (*Oryza sativa*), and maize (*Zea mays*) (Lurin et al., 2004; Schmitz-Linneweber and Small, 2008; Barkan and Small, 2014). Members of the PPR family are characterized by tandem arrays of a degenerate 35-amino-acid motif (PPR motif), and the PPR family is divided into P and PLS subfamilies, according to the nature of the PPR motifs. Members of the P subfamily function in various processes of RNA maturation in organelles, including RNA stabilization, splicing, intergenic RNA cleavage, and translation (Barkan and Small, 2014). The PLS subfamily contains canonical PPR (P) motifs, as well as long (L) and short (S) PPR-like motifs, in a P-L-S pattern. This subfamily is further divided into PLS, E/E+, and DYW classes, based on their C-terminal domains (Barkan and Small, 2014). PLS subfamily members function in RNA editing (Barkan and Small, 2014), a post-transcriptional modification of organelle transcripts, including C-to-U, U-to-C and A-to-I editing (Chateigner-Boutin and Small, 2010; Ruwe et al., 2013; Ruwe et al., 2019).

Kernel size and carbohydrate import into kernels directly determine the grain yield of maize, therefore, elucidation of the genetic basis of kernel traits could provide favorable alleles to enhance maize breeding. In a previous study, a maize recombinant inbred line (RIL) population was developed from a cross between two diverse parents, Zheng58 and SK (Small Kernel), which show dramatic variation in kernel weight; and a major kernel weight QTL, *qKW9*, was identified (Raihan et al., 2016; Yang et al., 2019). In this study, we mapped and cloned the causative gene underlying *qKW9,* and identified it as a PLS-DYW type PPR protein. We found that qKW9 is required for C-to-U editing at position 246 of *NdhB*, a chloroplast-encoded subunit of the NDH complex. Functional characterization revealed that C-to-U editing of *NdhB* is crucial for the accumulation of its protein product as well as the activity of the NDH complex. Impairment of this complex led to lower photosynthetic efficiency and a corresponding yield loss of maize in field trials.

## RESULTS

### Fine mapping and validation of *qKW9*

*qKW9* is a major QTL regulating maize kernel weight identified in the ZHENG58×SK RIL population (Raihan et al., 2016). Near-isogenic lines (NILs) harboring the *qKW9* allele from SK or ZHENG58 were screened from RIL-derived heterogeneous inbred families (HIFs) and used to fine map *qKW9*. In contrast to *dek* mutants, which have dramatic kernel weight loss due to defects in the embryo and/or endosperm, the NIL-SK kernels weighed only about 3g less per hundred kernels, compared to NIL-ZHENG58, and their kernel morphology, starch granule structure and plant morphology were similar (Figure 1, Table S1). Thus, the kernel development of NIL-SK plants was not strongly affected. Interestingly, two-week-old seedlings of NIL-SK were smaller than NIL-ZHENG58, possibly as a result of less nutrition from smaller kernels for their heterotrophic growth (Figure1B). However, at the mature stage, NIL-SK, and NIL-ZHENG58 plants had similar plant architecture(Figure 1A and 1C). NIL-SK plants had the same kernel row number but fewer kernels per row compared to NIL-ZHENG58 plants, resulting in smaller ears with fewer kernels (Figure 1D and Table 1).

**Figure 1.**
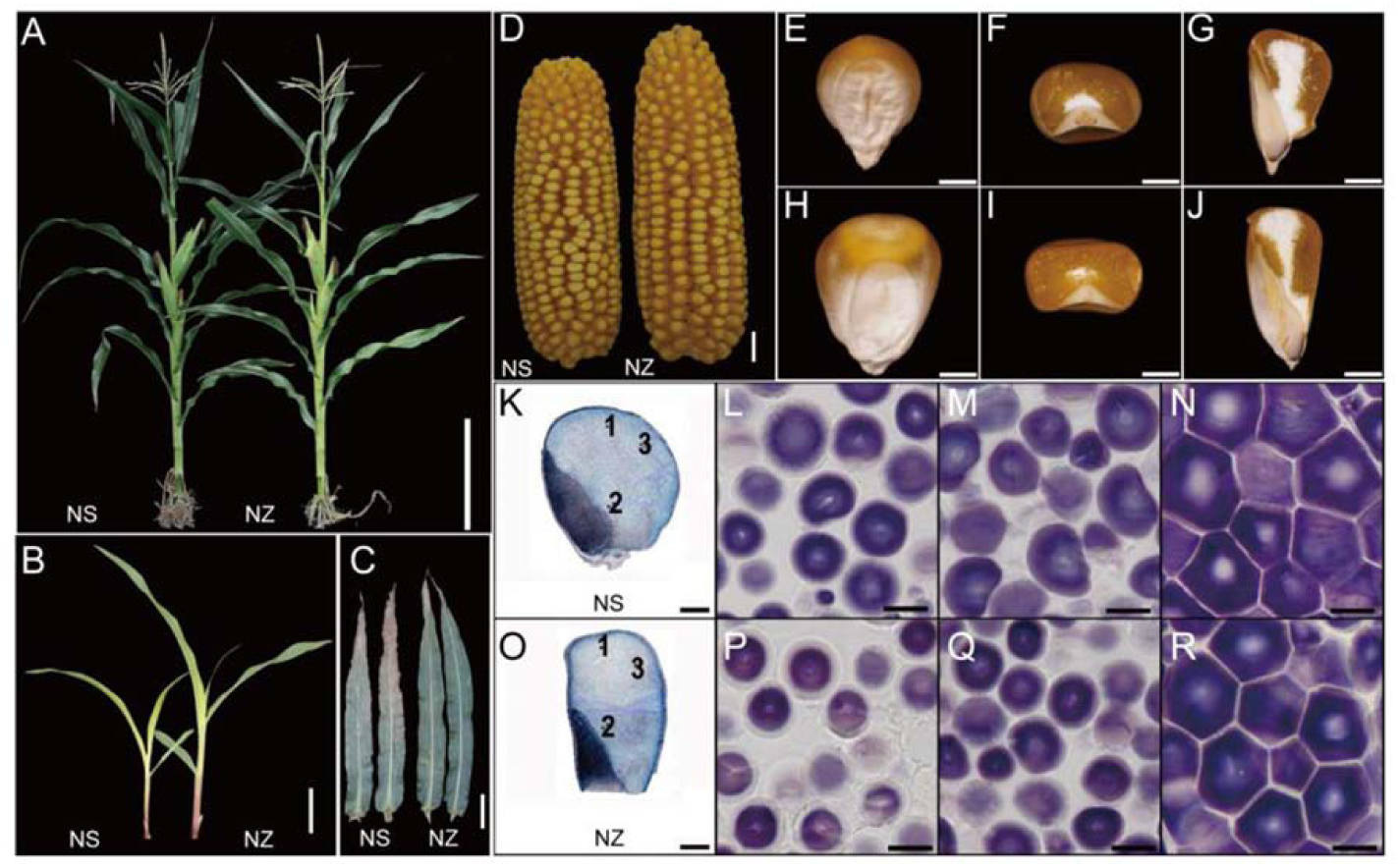
Plant and kernel morphology of NIL-SK and NIL-Zheng58. **(A)** NIL-SK (NS) and NIL-Zheng58 (NZ) had very similar plant architecture, Bar=20cm. **(B)** NIL-SK (NS) two-week old seedlings were smaller than NIL-Zheng58 (NZ). Bar=4cm. **(C)** Leaf senescence was greater in NIL-SK (NS) at 30 days after pollination compared to NIL-Zheng58. Bar=10cm. **(D)** Ears of NIL-SK (NS) were smaller than in NIL-Zheng58 (NZ). Bar=1cm. **(E) to (J)** Mature kernels of NIL-SK (E-G) were smaller that in NIL-Zheng58 (H-J). Whole kernels of NIL-SK and NIL-Zheng58 (**[E]** and **[H]**). Bar=2mm; transverse section of kernel of NIL-SK and NIL-Zheng58(**[F]** and **[I]**). Bar=2mm; Longitudinal section of kernel of NIL-SK and NIL-Zheng58(**[G]** and **[J]**). Bar=2mm; **(K) to (R)** Similar starch structure in endosperms of mature kernels of NIL-SK and NIL-Zheng58. Whole longitudinal section stained with iodine solution of kernels of NIL-SK and NIL-Zheng58 (**[K]** and **[O]**), 1, 2, 3 indicate the crown, farinaceous and keratin endosperm regions, respectively. Bar=1mm; **(L)** to **(N)** correspond to regions 1, 2, 3 in **(K)**; and **(P)** to **(R)** correspond to regions 1, 2, 3 in **(O)**. Bar=10μm.

**Table 1.** Ear related and agronomic traits in NIL-SK and NIL-Zheng58.

| Trait | NIL-SK |  | NIL-Zheng58 |  | P-value |
| --- | --- | --- | --- | --- | --- |
| | Mean ± SD<sup>a</sup> | N<sup>b</sup> | Mean ± SD | N | |
| Hundred Kernel Weight/g | 15.59 ± 2.01 | 27 | 18.57±1.21 | 30 | 6.07 × 10<sup>-9</sup> |
| Ear Length/cm | 9.75 ± 0.75 | 31 | 10.68±0.76 | 37 | 3.72 × 10<sup>-6</sup> |
| Kernel Number per Row | 22.00±2.99 | 27 | 24.03±2.28 | 32 | 4.47 × 10<sup>-3</sup> |
| Ear Row Number | 12.43±1.00 | 28 | 12.39±0.80 | 36 | 0.86 |
| Kernel Number per Ear | 249.22±39.45 | 27 | 279.39±34.28 | 31 | 2.89 × 10<sup>-3</sup> |
| Ear Weight/g | 43.79±7.60 | 30 | 57.33±8.35 | 36 | 3.74 × 10<sup>-9</sup> |
| Kernel Weight per Ear/g | 38.84±6.60 | 19 | 53.68±7.44 | 17 | 3.07 × 10<sup>-7</sup> |
| Plant Height/cm | 190.73±10.52 | 92 | 195.29±10.12 | 91 | 3.22 × 10<sup>-3</sup> |
| Ear Height/cm | 84.48±7.19 | 92 | 83.65±9.68 | 91 | 0.51 |
| Days to Shedding/Day | 63.38±1.81 | 68 | 60.16±1.21 | 70 | 6.72 × 10<sup>-24</sup> |

<sup>a</sup> SD=standard deviation; <sup>b</sup>N, number of observed individuals.

In previous study, line KQ9-HZAU-1341-1 from ZHENG58×SK RIL population with residual heterozygosity was used as founder line to fine map qKW9.2(Raihan et al., 2016; Liu et al., 2018). After three generations self-cross and screening against descendents of KQ9-HZAU-1341-1, several recombinant HIFs were obtained. Among the HIFs, F1H5 was used to generate recombination populations to screen for new recombinants to fine map *qKW9* in this study. Eight recombinants was identified by screening 685 F1H5 descendents and they were self-crossed for further analysis (Figure S1). By comparing Hundred Kernel Weight (HKW) of the homozygous progenies from all recombinants, *qKW9* was fine mapped to a ∼ 20kb region defined by markers M3484 and M3506 (156.65Mb and 156.67Mb, respectively in B73 RefGen v4) (Figure 2A and Fig S1). Three genes (*Zm00001d04850*, *Zm00001d048451*, and *Zm00001d048452*) were annotated within this region in B73 RefGen v4 (Figure 2B). Several SNPs were found in Zm00001d04850, however they were all synonymous. The second gene, Zm00001d048451, had a 13bp-CDS-deletion in SK, possibly leading to loss of function (Figure 2C). We failed to amplify the third gene, *Zm00001d048452*, from both SK and ZHENG58, and therefore, screened SK and ZHENG58 BAC libraries to search for sequence variation. However, sequence alignment and annotation revealed that *Zm00001d048452* was absent from both SK and ZHENG58, and there were no additional annotated genes within the *qKW9* locus, although there were some large-fragment insertions or deletions in the intergenic regions (Figure 2B). These results were further verified using the assembled SK genome (Yang et al., 2019). Of the two remaining candidates, Z*m00001d048450* displayed neither change in expression level nor pattern (Figure S2A), which together with its lack of non-synonymous SNPs suggested *Zm00001d048451* to be the causative gene of *qKW9*.

**Figure 2.**
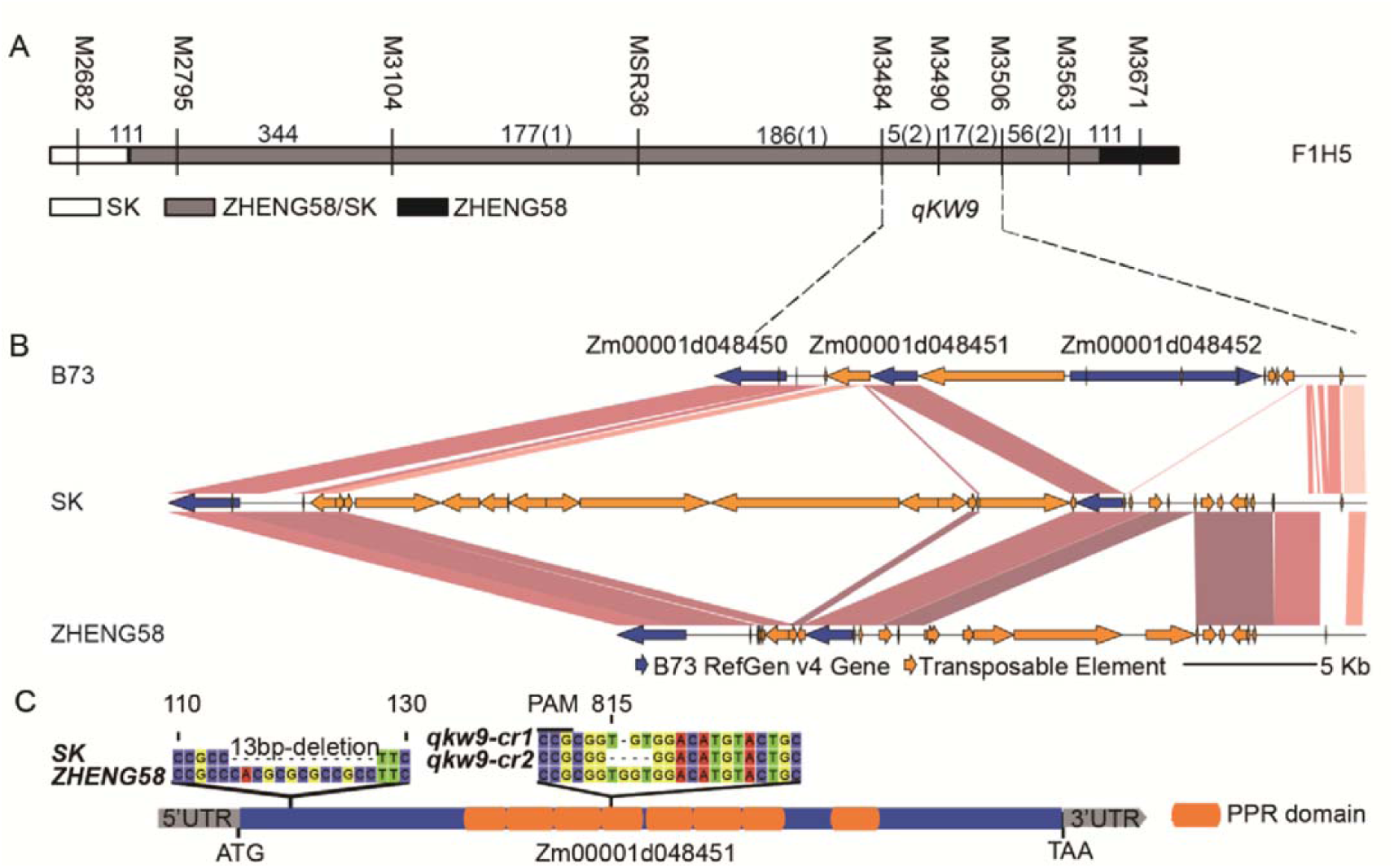
Fine mapping and gene structure of *qKW9*. **(A)** Mapping delimits *qKW9* to the region between M3484 and M3506 on chromosome 9. F1H5 which derives from ZHENG58 × SK RIL population is the founder line for screening heterozygous inbred families (HIFs) for fine mapping *qKW9*. Progeny tests of kernel weight were conducted on the resulting recombinant families. White bar represents the homozygous chromosomal segment for SK, grey bar represents the heterozygous chromosomal segment for ZHENG58 × SK, black bar represents the homozygous chromosomal segment for ZHENG58. The graphical genotype represents F1H5. Numbers between markers represent physical distances (Kb) between the adjacent markers and numbers in brackets represent the number of recombinants. **(B)** Gene annotations in the region of *qKW9* of B73, SK, and Zheng58. Sequences were obtained by sequencing BACs covering *qKW9* from SK and ZHENG58 genome BAC libraries, respectively. Zm00001d048452 was absent in both SK and ZHENG58. Two candidate genes-*Zm00001d048450* and *Zm00001d048451*-were identified in *qKW9*. **(C)** *Zm00001d048451* is a 1.8kb intron-less gene with 8 pentatricopeptide repeats and a 13bp-deletion was identified in coding region of *Zm00001d048451* in SK. CRISPR/Cas9 was used to create knockout mutants with a single guide sequence (the 20bp sequence adjacent to PAM) targeting *Zm00001d048451* in the inbred C01. Two mutated alleles - *qkw9-cr1* and *qkw9-cr2* - were identified by sequencing the first-generation (T_0_) plants and used for further genetic analysis.

To validate *Zm00001d048451* as the gene underlying *qKW9*, we adopted the CRISPR/Cas9 system to create knockout mutants (Figure 2C). Editing of *qKW9* was identified by Sanger sequencing of T_0_ transgenic plants, and two null mutants, *qKW9*-cr1, carrying a 1bp-deletion, and *qKW9*-cr2, carrying a 4bp-deletion, were used for subsequent analysis (Figure 2C and Figure 3). For both alleles, we found that kernel weight and ear weight were reduced compared to their corresponding wild type (Figure 3), demonstrating that Zm00001d048451 was indeed the causative gene of *qKW9*.

**Figure 3.**
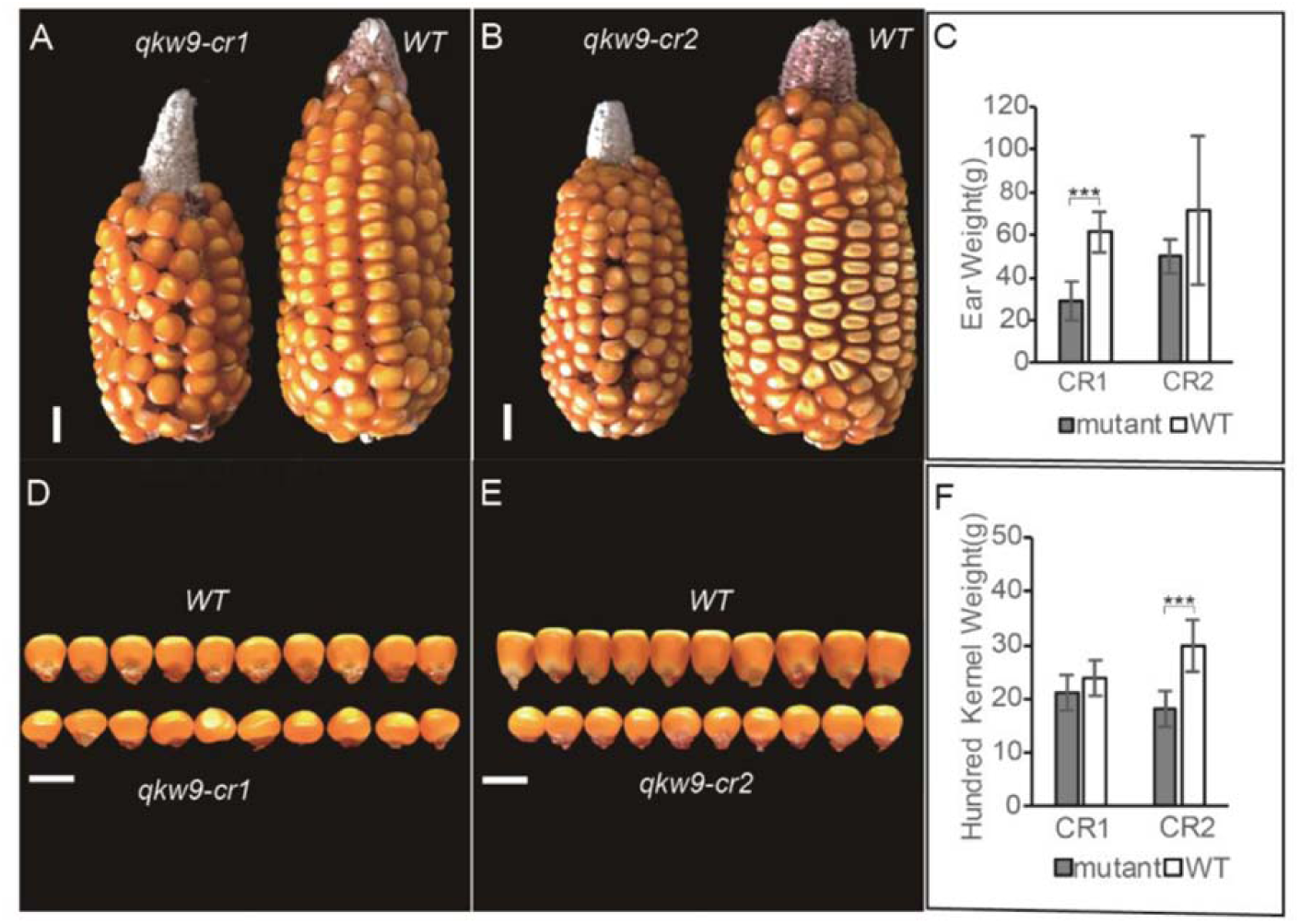
Two CRISPR/Cas9 knockout mutants of Zm00001d048451-*qkw9-cr1* and *qkw9-cr2*-produced smaller ears and smaller kernels than wild type. Each mutant is shown alongside its corresponding wild type segregant from a single Cas9-free T_1_ generation plant. **(A)** and **(B)** comparison of ears produced by CRISPR/Cas9 mutants (left) and WT (right). *qkw9-cr1* (left) and wild type (right) in **(A)** and *qkw9-cr2* (left) and wild type (right) in **(B)**. Bar=1cm. **(D)** and **(E)** kernels produced by CRISPR/Cas9 mutants (lower row) were smaller than WT (upper row). *qkw9-cr1* (lower) and wild type (upper) in **(D)** and *qkw9-cr2* (lower) and wild type (upper) in **(E)**. Bar=1cm. **(C)** and **(F)** show reductions in ear weight **(C)** and kernel weight **(F)** of CRISPR/Cas9 knockout mutants. Data are shown as mean ± SD (n=6). *** P < 0.001.

### *qKW9* is highly expressed in leaf, and encodes a chloroplast protein involved in *NdhB* RNA editing

*Zm00001d048451*/*qKW9* is predicted to encode a DYW subgroup pentatricopeptide repeat (PPR) protein with eight putative PPR motifs (Figure 2C). An Arabidopsis ortholog, *AT5G66520* (Figure 4A) encodes a DYW subgroup protein with ten PPR motifs and was designated *Chloroplast RNA Editing Factor 7* (*CREF7*), functioning in *Ndh* editing (Yagi et al., 2013). In order to address if *qKW9* is also involved in chloroplast RNA editing, we analyzed its expression and subcellular localization. Real-time PCR of *qKW9* revealed a considerably higher expression level in leaf than in other tissues (Figure S2B). *qKW9* expression was detected in all leaf-related tissues, and its expression level (13.9-87.6 FPKM) was much higher than in other tissues (0-12.8 FPKM) (Stelpflug et al., 2016). To test the subcellular localization of *qKW9*, we transiently expressed a *qKW9*-GFP fusion protein in tobacco, and found localization in the stroma of chloroplasts (Figure4B-4E), agreeing with a chloroplast prediction by TargetP (Emanuelsson et al., 2007).

**Figure 4.**
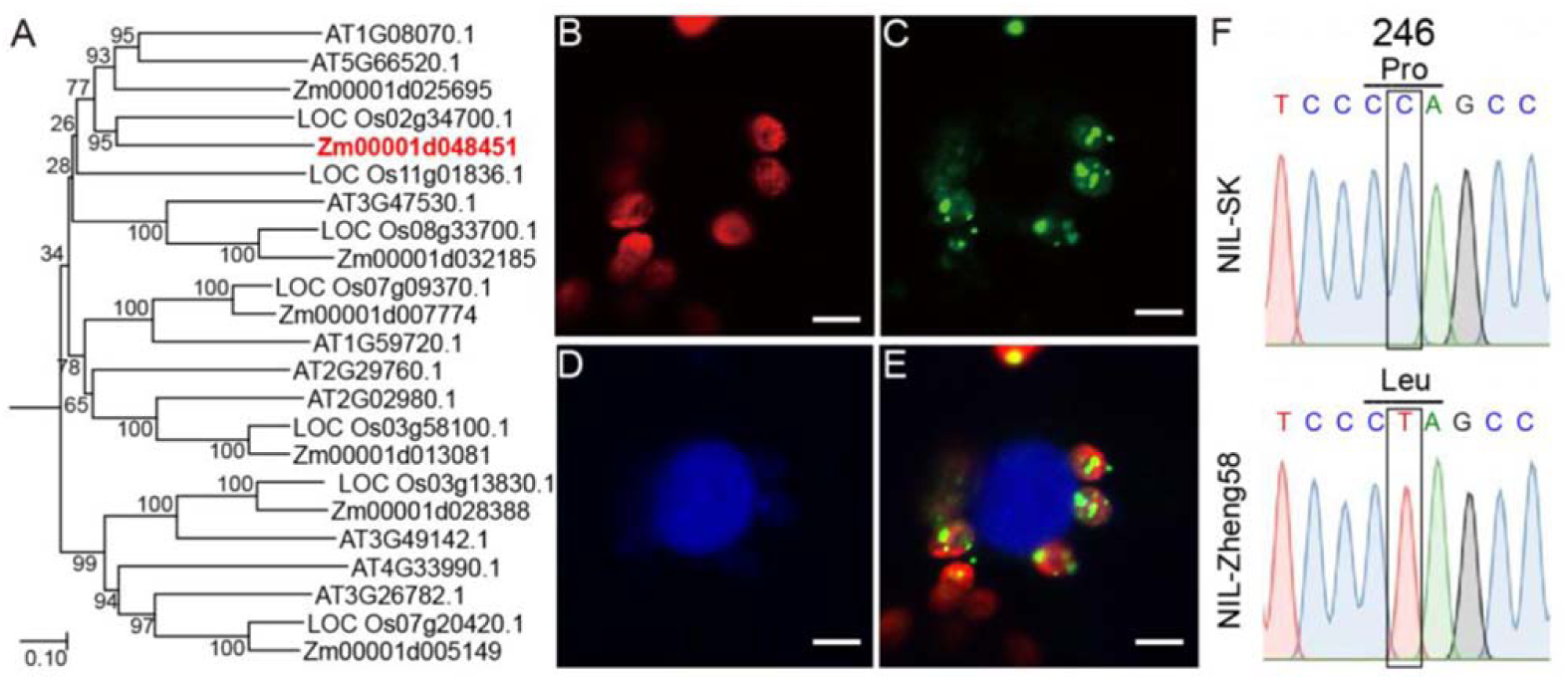
Characterization of *qKW9*/*Zm00001d048451*. **(A)** Phylogenetic tree of maize, Arabidopsis and rice PLS-E, PLE-E+, and PLS-DYW Pentatricopeptide Repeat genes predicted to localize in chloroplast/plastid by TargetP. **(B)** Autofluorescence of chlorophyll (red). **(C)** qKW9-GFP fusion protein (green) in green puncta within plastids **(D)** DAPI staining (blue) of nuclei **(E)** Overlay of **(B)**, **(C)** and **(D)**. Scale bar=5 µm. **(F)** Allele in NIL-SK of *Zm00001d048451* fails to edit C to U in 246^th^ codon of *NdhB* gene. C-to-U editing in NdhB-246 results in amino acid change from proline to leucine. Pro, proline; Leu, leucine.

To evaluate RNA editing by *qKW9*, leaves from NIL-SK and NIL-ZHENG58 plants before and after pollination were collected for total RNA sequencing. By comparing editing frequencies between NIL-SK and NIL-ZHENG58, six loci putatively edited by qKW9 were identified with *p*-value < 0.05 and mean editing frequency difference > 5% (Table 2). Three of these loci at chloroplast genome positions 90736, 132001, and 65407 (B73 RefGen v4), had striking editing differences between NIL-SK and NIL-ZHENG58, with close to 100% editing in NIL-ZHENG58 but almost none in NIL-SK at all stages tested (Table 2). Position 65407 is in an intergenic region, whereas positions 90736 and 132001 are in the 246^th^ codon of *GRMZM5G876106* and *GRMZM5G810298,* respectively (Table 2). These genes are the two copies of *NdhB* in the chloroplast genome, and their C-to-U editing changes the 246^th^ amino acid from proline to leucine. Thus, the sites edited by *qKW9* were designated as *NdhB-246*. We confirmed the *NdhB-246* editing difference by Sanger sequencing in NIL-SK and NIL-ZHENG58 (Figure 4F). We also investigated the editing frequency of *NdhB-246* in leaves of our two CRISPR/Cas9 null mutants, as expected, *NdhB-246* editing being abolished in both mutants (data not shown). These results demonstrate that *qKW9* is essential for *NdhB-246* editing.

**Table 2.**
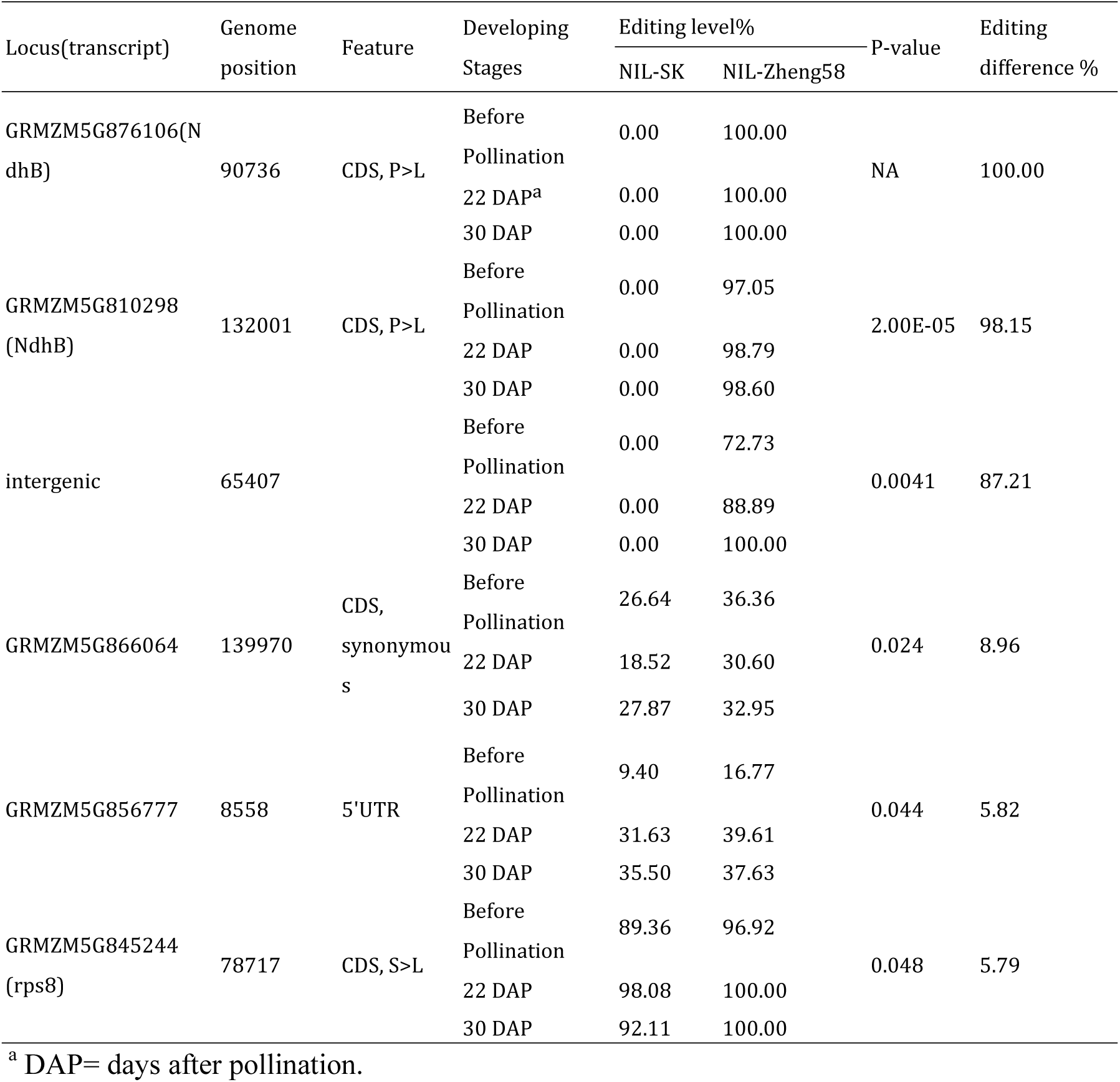
C-to-U editing sites in plastid genes with significant editing frequency difference between NIL-SK and NIL-Zheng58.

RNA editing defects may directly alter protein function or affect its ability to form complexes with other proteins (Hammani et al., 2009). *NdhB* encodes a subunit of the NDH complex (Laughlin et al., 2019), so we asked if this complex accumulates in the null *qKW9* background using protein blots probed with antibodies against NdhH to monitor accumulation of the complex. In NIL-SK, the level of NdhH was reduced to less than 25% of NIL-ZHENG58 (Figure 5), suggesting that *NdhB-246* RNA editing by *qKW9* is important for normal accumulation of the NDH complex.

**Figure 5.**
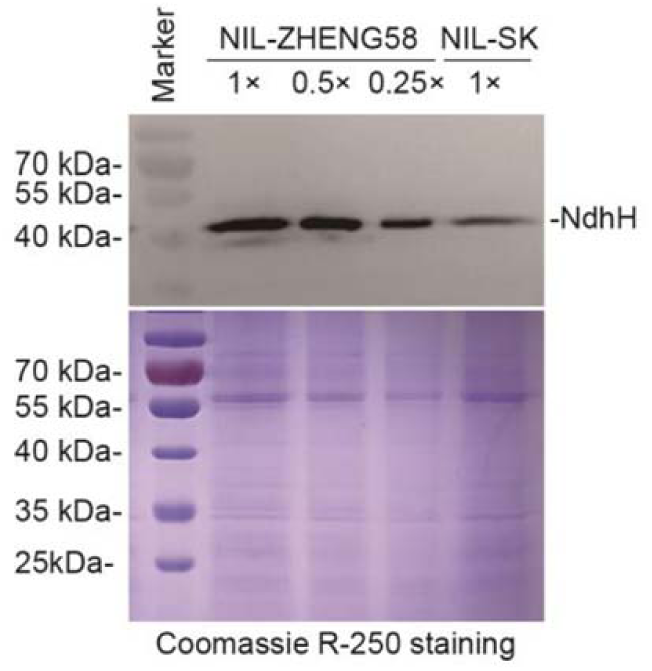
Protein blot analysis of the NDH complex. Chloroplast membrane protein was extracted with a commercial kit and protein samples were quantified with BCA protein assay. 1× sample amount equals 40 µg protein. Antibody against NdhH was used to indicate the amout of NDH complex. Chloroplast membrane protein from NIL-ZHENG58 was loaded a series of dilutions as indicated. Specific bands corresponded in size of NdhH protein (expected in 45 kDa, apparent in 49 kDa). Signals in NIL-ZHENG58 declined along with the dilution. The level of NdhH in NIL-SK was reduced to less than 25% of NIL-ZHENG58. Coomassie R-250 staining was used to show the proteins separated by electropheris as a loading control.

### C-to-U editing of *NdhB-246* is essential for optimal activity of NDH complex, electron transport rate and non-photochemical quenching induction

The chloroplast NADH dehydrogenase (NDH) complex transfers electrons originating from Photosystem I (PSI) to the plastoquinone pool, while concomitantly pumping protons across the thylakoid membrane (Strand et al., 2017). Its activity can be monitored as a transient increase in chlorophyll fluorescence, reflecting plastoquinone reduction after turning off actinic light (AL) (Burrows et al., 1998; Shikanai et al., 1998). In Arabidopsis, several nuclear mutants affecting NDH activity function in RNA processing of NDH subunit transcripts. For instance, *Chlororespiratory Reduction 2* (*CRR2*) functions in the intergenic processing of chloroplast RNA between *rps7* and *NdhB* (Hashimoto et al., 2003). A null allele of *CRR2* lacks NDH activity, and the post-illumination increase in chlorophyll fluorescence is undetectable, with a similar phenotype being observed in the tobacco *NdhB* mutant (Hashimoto et al., 2003).

To check whether *qKW9* impaired NDH activity, we monitered chlorophyll fluorescence using the post-illumination method (Burrows et al., 1998; Shikanai et al., 1998). Figure 6A shows a chlorophyll fluorescence trace from wild-type maize and *qkw9-cr1* and *qkw9-cr2*. In both *qkw9-cr1* and *qkw9-cr2*, the post-illumination increase of chlorophyll fluorescence was reduced, indicating that NDH activity was diminished in the null *qKW9* background, and that the Leu residue at position 246 of NdhB protein is required for NDH accumulation and activity. We next measured non-photochemical quenching (NPQ), a chlorophyll fluorescence parameter indicative of the level of thermal dissipation. NPQ was induced with increasing light intensity in both *qkw9*-cr1 and wild type prior to saturation of the ETR (Figure 6B). However, its induction in *qkw9*-cr1 was significantly lower at light intensities of 2413 μmols^-2^m^-1^, indicating that thermal dissipation was impaired in *qkw9*-cr1(Figure 6B). ETR represents the relative flow of electrons through PSII during steady-state photosynthesis. It increases with increases in light intensity until a point at which it cannot be further increased – termed its saturation point. For both wild-type and *qkw9*-cr1, the saturation point was over 400 μmol/m^2^s (Figure 6C). Whilst, ETR was not affected in *qkw9-cr1* at a low light intensities of ∼100 μmols-^2^m^-1^ (Figure 6C), it tended to be lower in *qkw9-cr1* at intensities above this (significantly so at 200 and 2400 μmols^-2^m^-1^). These altered ETR activities are consistent with the overall reduced grain yield in NIL-SK considering that light intensity is far in excess of 100 μmols-^2^m^-1^ in the field. These findings considerably differ from studies in Arabidopsis and tobacco, since multiple mutant analyses have demonstrated that NDH does not function to trigger thermal dissipation in PSII in these species (Burrows et al., 1998; Munekage et al., 2004).

**Figure 6.**
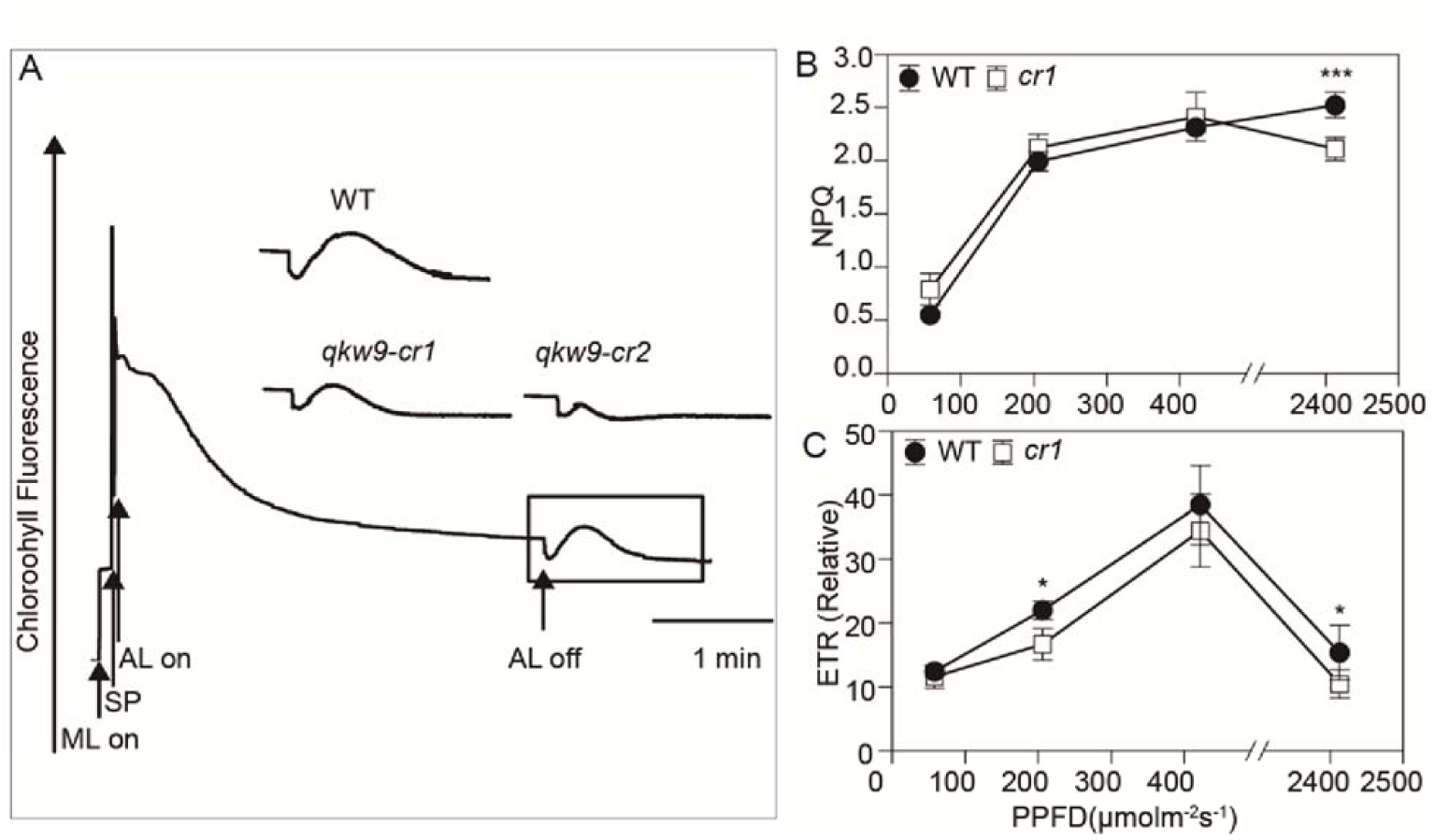
NDH activity monitoring and NPQ and ETR in null mutants of qKW9. **(A)** Monitoring of NDH activity using chlorophyll fluorescence analysis for *qkw9-cr1* and *qkw9-cr2* mutants. The curve shows the typical change trace of chlorophyll fluorescence *in vivo* as the NDH complex catalyzes the post-illumination reduction of the plastoquinone pool (Okuda et al., 2007). The change in post-illumination fluorescence ascribed to NDH activity was different between WT and mutants. Insets are magnified traces from the boxed area. ML, measuring light; AL, actinic light; SP, a saturating pulse of white light. **(B)** NPQ was induced by light intensity in both *cr*1 and WT, but it was significantly lower in *cr*1 under photon flux density of 2413 μmol of photons m^-2^s^-1^. **(C)** relative ETR (rETR) under different photon flux densities. rETR in cr1 and WT reached maximum when the light intensity was 422 μmol of photons m^-2^s^-1^. It was significantly lower in *cr*1 under the photon flux density of 206 μmol of photons m^-2^s^-1^ and 2413 μmol of photons m^-2^s^-1^. The rETR is depicted relative to a maximal value of ϕ_PSII_ × PPFD (photon flux density, μmol of photons m^-2^s^-1^). Data are shown as mean ± SD (*n*=6).

### *qKW9* controls kernel weight by regulating photosynthesis

Genetic evidence suggests that physiological functions of cyclic electron transport (CET) around Photosystem I (PSI) are essential for efficient photosynthesis and plant growth (Munekage et al., 2004). The physiological role of CET is to protect PSII under intense light via ΔpH-dependent thermal dissipation in PSII, as well as to act as an ATP generator in photosynthesis (DalCorso et al., 2008; Alric and Johnson, 2017). Our results suggest that reduced activity of the NDH complex in maize affected NPQ and ETR. We therefore asked how photosynthesis and carbon assimilation were affected by changes in NDH activity? We measured the fresh weight of developing kernels of NIL-SK and NIL-ZHENG58 under field conditions, and investigated several photosynthesis-related parameters (Figure 7). Fresh weight of NIL-SK kernels was similar to NIL-ZHENG58 before 30 DAP (Figure 7A). However, kernels of NIL-SK reached their maximum fresh weight at 30DAP, while NIL-ZHENG58 kernels continued to gain weight until 35 DAP, suggesting that carbon deposition in kernels was greater in NIL-ZHENG58 at 35 DAP (Figure 7A). Consistent with this observation, leaves of NIL-SK had more severe senescence at 30 DAP compared to NIL-ZHENG58, indicating decreased source strength in the NIL-SK plants (Figure 1C). NIL-SK also had significantly lower net photosynthesis than NIL-ZHENG58 at 22DAP and 30DAP (Figure 7B). Consistently, stomatal conductance and transpiration rate were similarly lower in NIL-SK than in NIL-ZHENG58 (Figure 7C-7D). The lower photosynthetic capacity of NIL-SK, coupled with the potential compensatory fact that less kernels were produced per ear in this line (Table 1), may explain why the fresh weight of NIL-SK were not significantly lower than NIL-ZHENG58 at 22DAP and 30DAP (Figure 7A). In addition, the chlorophyll content (SPAD value) and the maximum photochemical efficiency (Fv/Fm) were invariant between the NILs (Figure 7E-7F), indicating that the differences in the net photosynthetic rates might not result from a different level of photosynthesis potential. Accordingly, the photosynthetic rate was also significantly lower in *qKW9*-cr1 than WT at 30DAP under field conditions (*qKW9*-cr1: 17.35±2.10 μmol CO_2_m^-2^s^-1^, Wild type: 29.68±3.56μmol CO_2_ m^-2^s^-1^, *n*=6). We conclude that impaired NDH activity affected both net photosynthesis and the duration of active photosynthesis, resulting in smaller ears and kernels in NIL-SK.

**Figure 7.**
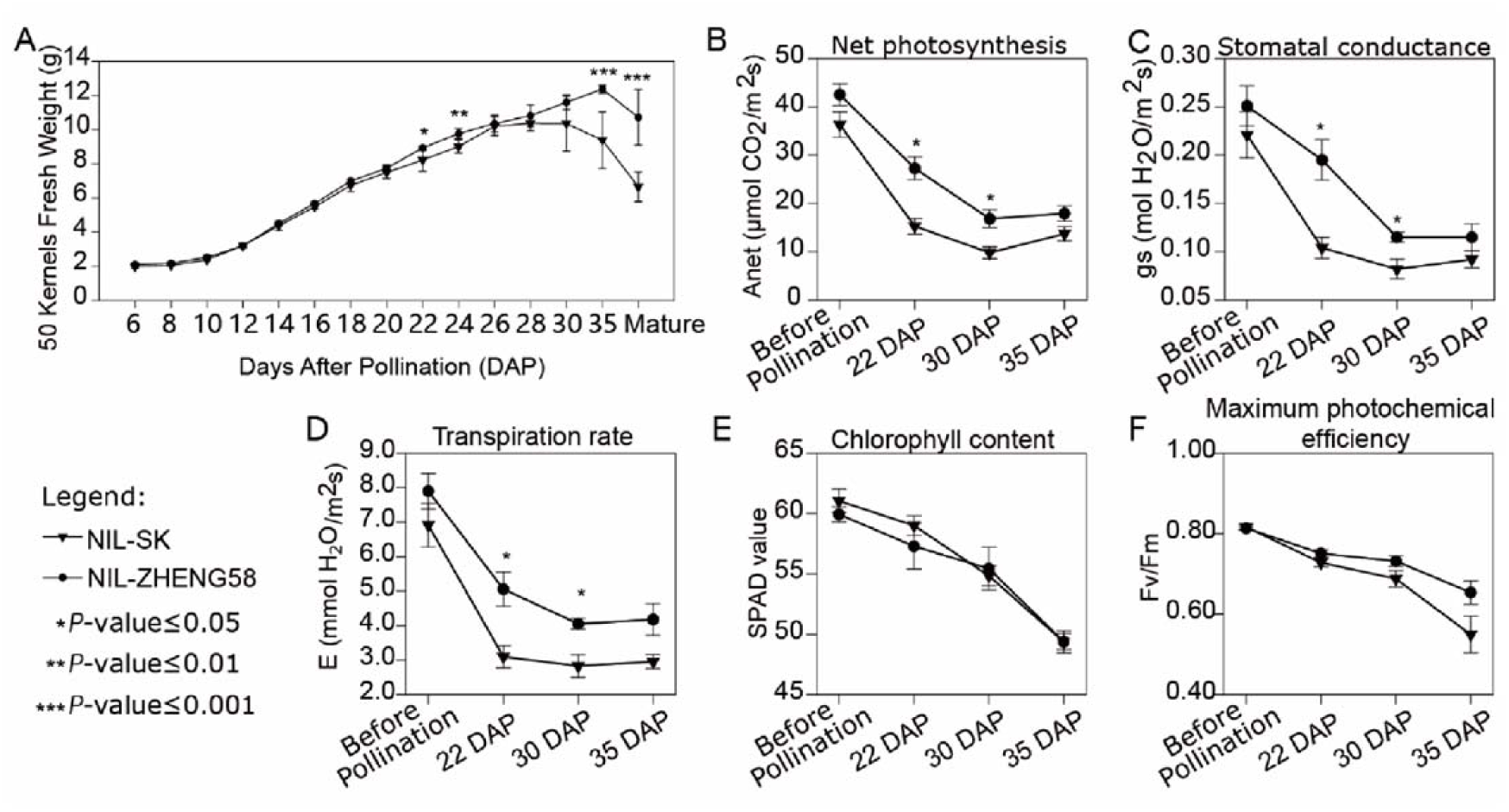
Grain filling and photosynthesis measurement in NIL-SK and NIL-Zheng58. **(A)** Time courses of fresh weight of 50 kernels of NIL-SK and NIL-Zheng58. The fresh weight of NIL-SK and NIL-ZHEGN58 reached the maximum at 30 DAP and 35 DAP, respectively. **(B)** to **(E)** Time courses of photosynthesis-rate related parameters of NIL-SK and NIL-Zheng58. Net photosynthesis **(B)**, stomatal conductance **(C)**, and transpiration rate **(D)** were significantly lower in NIL-SK than NIL-ZHENG58 at 22 DAP and 30 DAP; **(E)** chlorophyll content and **(F)** maximum photochemical efficiency did not show significant between genotype differences at any of the four stages tested. Data are shown as mean ± SD (*n*=6).

## DISCUSSION

### *qKW9* encodes a PPR gene responsible for C-to-U editing of NdhB

The maize kernel has been of interest to researchers as a model system for the study of development and genetics for a century. Numerous kernel mutants have been identified (Neuffer and Sheridan, 1980; Sheridan and Neuffer, 1980; Clark and Sheridan, 1991), and in recent years, many mutants that result in dramatically reduced kernel size, and seedling lethality have been identified. In many cases, *PPR* genes are responsible for these phenotypes, due to their function in organellar gene expression. Generally, null alleles of PPR genes in previous studies produce kernels that are with obvious development abnormality at early stages and are easily distinguished from normal kernels on self-crossed F_1_ ears due to their smaller size, pale pericarp, flat or shrunken appearance (Manavski et al., 2012; Sosso et al., 2012; Li et al., 2014; Sun et al., 2015; Xiu et al., 2016; Cai et al., 2017; Chen et al., 2017; Qi et al., 2017; Ren et al., 2017; Dai et al., 2018; Li et al., 2018; Sun et al., 2019). Unlike these kernel mutants, kernels produced by null allele of *qKW9* are similar in appearance and viability although smaller in size comparing to wild type, and kernel weight is determined by genotype of maternal plant rather than kernel genotype. *qKW9* is the first identified C-to-U editing factor in the maize chloroplast that has a quantitative rather than qualitative effect on kernel and ear size. This difference stems from the involvement of *qKW9* in the abundance of the NDH complex, which in known to play a regulatory role in photosynthesis (Nashilevitz et al., 2010; Peltier et al., 2016). Based on results from this study, it is possible that variants of other yet-unidentified RNA editing factors responsible for the 11 C-to-U editing sites in maize *ndh* transcripts will also affect kernel and ear size in a quantitative way. Indeed, studies in *Arabidopsis* have identified many PPR genes affecting ndh expression, by focusing on changes in chlorophyll fluorescence related to NDH activity (Kotera et al., 2005; Okuda et al., 2007; Hammani et al., 2009; Okuda et al., 2010). Similar research in maize may help us rapidly identify potential QTL, circumventing the tedious work of fine mapping.

Grains are typical sink organs -i.e. they are net receivers of photoassimilates from photosynthetically active source tissues - and a considerable number of studies suggest that enhancing photosynthetic efficiency could increase the productivity of crops (Sonnewald and Fernie, 2018; South et al., 2019; Wu et al., 2019). In the light reactions of photosynthesis, linear electron transport (LET) from water to NADP^+^ does not fully satisfy the ATP/NADPH production ratio required by the Calvin-Benson cycle and photorespiration (Yin and Struik, 2018). Cyclic electron transport (CET) around photosystem I (PSI) has, therefore, been considered as a candidate for augmented ATP synthesis in response to fluctuating demand during photosynthesis (Rumeau et al., 2007; DalCorso et al., 2008; Nakamura et al., 2013). In PSI CET, electrons are recycled around PSI generating ΔpH and consequently ATP without a concomitant accumulation of NADPH (Shikanai, 2007; Munekage and Taniguchi, 2016). In Arabidopsis, two CET pathways have been identified by genetics. The main pathway depends on PROTON GRADIENT REGULATION 5 (PGR5)/PGR5-LIKE PHOTOSYNTHETIC PHENOTYPE 1 (PGRL1) proteins, whereas the minor pathway is mediated by the chloroplast NADH dehydrogenase-like (NDH) complex (Munekage et al., 2004). The Arabidopsis *pgr5* mutant is defective in PSI CET and was discovered by screening for reduced non-photochemical quenching (NPQ) of chlorophyll fluorescence (Munekage et al., 2002; DalCorso et al., 2008). However, as in tobacco, the Arabidopsis *chlororespiratory reduction* (*crr*) mutants and *organelle transcript processing* (*otp*)mutants defective in chloroplast NDH did not show any marked phenotype (Burrows et al., 1998; Hashimoto et al., 2003; Munekage et al., 2004; Hammani et al., 2009; Okuda et al., 2010).

Plastid genomes encode 11 subunits (NdhA to NdhK) forming the core of the membrane arm of the L-shaped structure of the NDH complex (Laughlin et al., 2019). Multiple PPR genes regulate expression of *ndh* genes in Arabidopsis. These PPR genes are either responsible for splicing of polycistronic transcripts, or site-specific C-to-U RNA editing (Hashimoto et al., 2003; Munekage et al., 2004; Kotera et al., 2005; Okuda et al., 2007; Hammani et al., 2009; Okuda et al., 2009; Okuda et al., 2010). C-to-U RNA editing is important in organelle gene expression in various organisms, although the efficiency varies in different organs and at different developmental stages (Maier et al., 1995; Peeters and Hanson, 2002). C-to-U RNA editing in Arabidopsis can generate translational initiation codons, as in *CRR4* (Kotera et al., 2005) or cause amino acid alterations, as in *CRR21*, *CRR22*, *CRR28*, *OTP82*, *OTP84* and *OTP85* (Okuda et al., 2007; Hammani et al., 2009; Okuda et al., 2009; Okuda et al., 2010). Editing of *NdhB-246* in leaf tissues is near 100% in maize, suggesting that it is important for the function of NdhB protein (Peeters and Hanson, 2002). *NdhB-246* editing also occurs in tobacco and rice (Tsudzuki et al., 2001). Therefore, C-to-U editing of *NdhB-246* appears crucial to its function. The *qKW9* QTL characterized in our study is the first RNA editing factor that has been linked to C-to-U editing of *NdhB-246*. Our results clearly indicate that the abolition of C-to-U editing in *NdhB-246* impairs accumulation of the NDH complex *in vivo*.

### Cyclic electron transport via NDH complex in C_4_ and C_3_

Chloroplast NDH mediation of CET around PSI was first reported in tobacco following disruption of *NdhB* (Shikanai et al., 1998). Knockout lines of *ndh* genes were created by plastid transformation in tobacco. Mutants defective in expression of chloroplast *ndh* genes were isolated in Arabidopsis (Hashimoto et al., 2003). However, both knockout lines of *ndh* genes in tobacco and mutants with impaired NDH activity in Arabidopsis lack morphological phenotypes (Hashimoto et al., 2003; Munekage et al., 2004; Okuda et al., 2007; Hammani et al., 2009; Okuda et al., 2009; Okuda et al., 2010). So, the general conclusion has been that mutants defective in NDH do not show a clear phenotype and NDH is dispensable at least when plants are grown in controlled environments. Analysis of this observation leads to the conclusion that NDH does not function to generate a pH gradient in order to trigger thermal dissipation in PSII. The observations we present here, where we found that *qKW9* is required for the expression of *NdhB* and optimal activity of the NDH complex in maize, suggest that this conclusion may not hold in all species. Namely in the null mutant genotypes, light intensity-dependent ETR and NPQ were obviously affected. We additionally observed more severe leaf senescence in NIL-SK, a phenonomon that may result from impaired thermal dissipation, since NPQ induction under high-intensity light is suppressed as compared with NIL-ZHENG58. Moreover, the overall rate of photosynthesis was also reduced in the null *qKW9 mutant*, leading to significantly reduced ear and kernel size.

Our study of *qKW9* provides a possible explanation for the apparent contradiction of these observations, in suggesting that CET via the NDH complex is more important in C_4_ than in C_3_ plants. CET around photosystem I is critical for balancing the photosynthetic energy budget of the chloroplast by generating ATP without net production of NADPH (Ishikawa et al., 2016a). C_4_ plants have higher ATP requirements than C_3_ plants (Ishikawa et al., 2016b), rendering the ATP supply by CET particularly important, imply that during the evolution of NADP-malic enzyme-type C_4_ photosynthesis in the C_4_-like genus *Flaveria*, CET was promoted by markedly increasing expression of both PGR5/PGRL1 and NDH subunits (Nakamura et al., 2013). The NDH subunit, however, increases markedly in bundle sheath cells with the activity of the C_4_ cycle while PGR5/PGRL1 increases in both mesophyll and bundle sheath cells in *Flaveria* and other C_4_ species, implying that the NDH complex provides a considerable role in the establishment of C_4_ photosynthesis (Nakamura et al., 2013). Previously, it was also shown that NDH plays a central role in driving the CO_2_-concentrating mechanism in C_4_ photosynthesis (Takabayashi et al., 2005; Andrews, 2010). In addition, the NDH complex has been experimentally demonstrated to be a high-efficiency proton pump, increasing ATP production by cyclic electron transport (Strand et al., 2017). Ishikawa et al. report that NDH-suppressed C_4_ plants are characterized by consistently decreased CO_2_ assimilation rates, impaired proton translocation across the thylakoid membrane and reduced growth rates (Ishikawa et al., 2016a). Results from our study provide direct evidence that the NDH complex is important to C_4_ photosynthesis. As such comparison of our data with that from a recent study in Arabidopsis which showed that NDH-dependent cyclic electron transport around PSI contributes to the generation of proton motive force only in the weak mutant of *pgr5* (Nakano et al., 2019), suggests that it is more important to C_4_ than C_3_ photosynthesis.

An alternative explanation could be the difference in the study conditions. In our study, growth and photosynthesis measurement are conducted in the field where the plants experience naturally fluctuating light, humidity, temperature, etc. Fluctuating elements may give rise to stress to photosystems. Impairment of NDH-dependent PSI cyclic electron transport causes a reduction in photosynthetic rate under fluctuating light, leading to photoinhibition at PSI and consequently to reductions in plant biomass in rice (Yamori et al., 2016). It is worth noting that there are studies stressing the role that NDH complex plays in CET under stresses (Horváth et al., 2000; Wang et al., 2006; Yamori et al., 2015). Indeed the conclusions that the NDH complexs role in CET is dispensable come from studies of tobacco and Arabidopsis grown in growth chambers rather than the field, resulting in an under-estimation of the importance of the NDH complex’s regulatory effect on photosynthesis.

In summary, our study identified a maize kernel size QTL which is caused by allelic variation in *qKW9,* a PLS-DYW type PPR protein. We found that qKW9 is required for C-to-U editing at 246^th^ codon of *NdhB*, a chloroplast-encoded subunit of the NDH complex. With this editing pattern previously being recorded occuring concomitantly with the onset of photosynthetic activity in tobacco (Karcher and Bock, 2002). Functional characterization revealed that C-to-U editing of maize *NdhB* is crucial for the accumulation of its protein product as well as the activity of the NDH complex. This study thus challenges current models of the role of the NDH in photosynthesis, revealing new insights into the regulation of C_4_ photosynthesis and suggesting a novel potential target for crop improvement.

## METHODS AND MATERIALS

### Fine mapping of *qKW9*

Multiple major QTL regulating kernel-size-related traits were identified by multi-environment QTL analysis in ZHENG58×SK RIL population and a major QTL on chromosome 9 regulating kernel width was designated as *qKW9* in a previous report (Raihan et al., 2016). To fine-map *qKW9*, the heterogeneous inbred family (HIF) was screened against the RIL population and RIL line KQ9-HZAU-1341-1 was heterozygous between Marker M2682 (155.83Mb in B73 Ref Gen v4) and Marker M3671 (156.83Mb in B73 RefGen v4) was used as the founder HIF (Raihan et al., 2016). In a nursery grown in Hainan in 2015, two groups of homozygous progenies of F1H5, which was a descendant of line KQ9-HZAU-1341-1, were significantly different in hundred kernel weight (HKW), kernel length (KL), and kernel width (KW). Thus, F1H5 was used as the starting HIF to fine map *qKW9* in this study. In the summer of 2016, recombinants between Marker M2795 and Marker M3671 were screened against the F1H5 population. In the winter of 2016, progeny tests were conducted on those recombinant populations.

For genotyping, genomic DNA extraction from young leaf was conducted using the CTAB protocol for plant tissues. To detect SNP and Indel markers, PCR was conducted in 10 μL reactions with KASP master mix (cat no: KBS-1030-002, LGC), self-made KASP array mix, and DNA template in 96 well non-transparent plates. KASP array mix was made by mixing equal volumes of primer F1 (36 μM), F2 (36 μM), and R (90 μM) of a specific SNP marker. For each reaction, 0.14 μL array mix, 1×master mix, and 20∼200 ng DNA were used. Thermal cycling was 94 °C for 15 minutes to activate the enzyme, followed by 10 cycles of touch down PCR (denature at 94 °C for 20 s, annealing/elongation start with 61 °C for 60 s, drop 0.6 °C per cycle), then annealing/elongation for another 26-36 cycles depending on the quality of primers (denature at 94 °C for 20 s, annealing/elongation at 55 °C for 60 s). Upon the completion of the KASP PCR, reaction plates were read by CFX96 Touch^TM^ Real-time PCR detection system and the data was then analysed with the Allelic Discrimination module of BioRad CFX Manager 3.0. Detected signals were plotted against FAM and HEX fluorescence intensity as a graph, with samples of the same genotype clustering together. To detect SSR markers, PCR products were detected by AATI Fragment Analyzer following the manufacturer’s instructions.

Maize plants were examined under natural field conditions in the experimental fields of Wuhan (30°N, 114°E), Sanya (18°N, 109°E), and Baoding (39°N, 115°E) in China. The planting density was 25 cm between adjacent plants in a row and the rows were 60 cm apart. Field management, including irrigation, fertilizer application, and pest control, essentially followed the normal agricultural practices. Harvested maize ears were air-dried and then fully-developed ears were shelled for measuring HKW, KL, and KW as previously reported (Raihan et al., 2016).

### BAC screen, sequence, and *de novo* assembly

BACs covering *qKW9* of both parent lines-SK and ZHENG58-were screened. BAC DNA was prepared using the QIAGEN Large-Construct Kit (Cat no: 12462) following the manufacturer’s instructions but with 150ml overnight-cultured bacterial input. The recovered DNA was sent to a company (Nextomics Bioscience Co., Ltd, Wuhan, China) for quality control and library construction. The resulting sequence data was assembled by PacBio’s open-source SMRT Analysis software.

### Fresh weight during the filling stage

NILs derived from homozygous progenies of HIF-p11 were used to analyze the grain filling rate of developing kernels after pollination. NILs with the *qKW9* allele of SK designated as NIL-SK while NILs with the *qKW9* allele of ZHENG58 designated as NIL-ZHENG58. Starting from 6 days after pollination (DAP), 50 fresh kernels were harvested and weighted from 6 ears of each NIL every other day until 30 DAP. At 35 DAP and upon harvest fresh kernels were also weighed.

### Mutagenesis of *qKW9* with CRISPR/Cas9-based gene editing

Two guide RNA sequences (cggtggtggacatgtactg and ctgttctggggatccagct) against *qKW9* were designed by CRISPR-P 2.0 (http://crispr.hzau.edu.cn/CRISPR2/) then cloned into a CRISPR/Cas9 plant expression vector (Liu et al., 2017a). The backbone of the vector was provided by WIMI Biotechnology Co., Ltd (Changzhou, China). The vector allows expression of single guide RNA by the ZmU61 promoter and Cas9 by a maize UBI promoter. The resulting binary plasmids were transformed into the Agrobacterium tumefaciens strain EHA105 and used to transform maize inbred C01. All constructs were sequence-verified.

### Light Microscopy

Whole sections of mature kernels were stained with iodine solution using the method in a previous report (Zhao et al., 2016). Three different regions of endosperm were examined for the morphology of starch.

### Subcellular localization of *qKW9*

*Zm00001d048451* was predicted to locate in chloroplast by TargetP (Emanuelsson et al., 2007). To verify this, a codon-optimized CDS (optimized by a web tool - https://www.genscript.com/codon_opt_pr.html) was fused with green fluorescent protein (GFP) and driven by expression from the cauliflowner mosaic virus 35S promoter. The binary vector-pK7FWG2.0-was obtained from Dr. Hannes Claeys (Cold Spring Harbor Laboratory, USA). The plasmids containing the chimeric genes were transferred into Agrobacterium tumefaciens strain GV3101. The resulting strain was co-infiltrated into *Nicotiana benthemiana* leaves with a strain harboring P19 which was obtained from Dr. Edgar Demesa Arevalo (Cold Spring Harbor Laboratory, USA) (Lindbo, 2007). Fluorescence signals were detected using LSM780. DAPI (4,6-diamidino-2-phenylindole) staining solution (http://cshprotocols.cshlp.org/content/2007/1/pdb.rec10850.full?text_only=true) was injected to the leaf before observing the fluorescence signals. Agrobacterium growth and injection followed the steps described in a previous report (Xu et al., 2015).

### Phylogenetic analysis

To identify the PPR genes in maize B73 RefGen v4, protein sequences of B73 RefGen v4 genes were downloaded from ftp://ftp.gramene.org/pub/gramene/ (B73 RefGen v4.59). Then HMMER 3.0 software (Finn et al., 2011) was used to scan all of the annotated Pentatricopeptide Repeat genes in B73 RefGen v4 with the Midden Markov (HMM) profile of PPR domain (PF01535.20, http://pfam.sanger.ac.uk/) (E-value < 1). Based on the C-terminal domain structure, the HMM profiles of E, E+, and DYW domain were rebuilt using the previously predicted PPR genes in B73 RefGen v3. Then these HMM profiles were used to scan the PPR genes annotated in B73 RefGen v4 (ftp://ftp.gramene.org/pub/gramene/CURRENT_RELEASE/gff3/zea_mays/gene_function). Then TargetP version 1.1 (http://www.cbs.dtu.dk/services/TargetP/) was used to predict the organelle targeting of these E, E+, and DYW types PPR proteins. Only the chloroplast targeting genes were kept to conduct the evolutionary analysis with their orthologous genes in Arabidopsis and rice (https://download.maizegdb.org/Zm-B73-REFERENCE-GRAMENE-4.0/Orthologs/). The evolutionary history was inferred using the Neighbor-Joining method (Saitou and Nei, 1987) by MEGAX (Kumar et al., 2018). The bootstrap consensus tree inferred from 500 replicates was taken to represent the evolutionary history of the taxa analyzed (Felsenstein, 1985).

### Photosynthetic parameters and chlorophyll content measurements

Carbon dioxide assimilation rate, stomatal conductance, and transpiration rate were measured on fully-expanded maize leaves grown in the field using a portable gas exchange system (LI-6400XT, LI-COR Inc., USA) as described (Huang et al., 2009; Bihmidine et al., 2013). The measurements were conducted at an ambient CO_2_ concentration of 400 μmol mol^−1^ and light saturation of 2000 μmol m^−2^ s^−1^. Leaf photochemical efficiency (Fv/Fm) was measured on dark-adapted leaves using the FlourPen FP100 chlorophyll fluorescence meter (Photon System Instruments, Czech Republic). Leaf chlorophyll content was measured using a chlorophyll meter (SPAD-502, Konica Minolta, Japan). The measurements were performed before pollination and at 22, 30, and 35 days after pollination (DAP) on eight NIL-SK plants or NIL-ZHENG58 plants. Means and standard errors (SE) were calculated using Microsoft Excel. Differences in chlorophyll content and photosynthetic parameters were assessed using the Student’s *t*-test embedded in the Microsoft Excel program, at the *P*-value ≤ 0.05 level.

### RNA sequencing

To explore the possible RNA editing in leaf by the PPR gene, the ear leaves from NIL-SK and NIL-ZHENG58 plants before pollination and after pollination (22 days and 30 days) were collected. Total RNA was isolated from these samples using Direct-zol RNA MiniPrep Plus kit (Cat no: R2072, ZYMO Research, USA). Libraries were constructed using the Illumina TruSeq Stranded RNA Kit (Illumina, San Diego, CA, USA) following the manufacturer’s recommendations. Strand-specific sequencing was performed on the Illumina HiSeq 2000 system (paired-end 100-bp reads). The raw reads were trimmed by Trimmomatic v0.36 (Bolger et al., 2014) to gain high-quality clean reads, and the quality of the clean reads was checked using the FASTQC program (Andrews, 2010). Next the clean reads were aligned to maize B73 RefGen v4 chloroplast genome by Hisat2 (Kim et al., 2015). Picard tools were subsequently used to add read groups, sort, mark duplicates, and create index (http://broadinstitute.github.io/picard/). Then the GATK was used to call the sequence variants by HaplotypeCaller (McKenna et al., 2010). The ratio of edited allele reads count/total reads count served as editing frequency for each site and the significance of editing frequency difference between NIL-SK and NIL-ZHENG58 was estimated by pairwise *t*-test with a threshold *P*-value < 0.05. Only the loci with a mean editing frequency difference over 5% between NIL-SK and NIL-ZHENG58 were treated as possibly RNA editing sites being affected by *qKW9*.

### RNA extraction and qRT-PCR

Total RNA was extracted from various plant tissues except leaf using RNA extraction kit (Cat no: 0416-50, Huayueyang, China). cDNA was synthesized from the extracted RNA using the TransScript One-Step gDNA Removal and cDNA Synthesis SuperMix (Cat no: AT311, TransGen Biotech, China). qRT-PCR was carried out in a total volume of 20 μl containing 2 μl of 10x-diluted reverse-transcribed product, 0.2 mM gene-specific primers, and 10 μl KAPA SYBR® FAST qPCR Master Mix (Cat no: KK4607), using a Bio-Rad CFX96 Touch^TM^ Real-time PCR detection system according to the manufacturer’s instructions. Quantitative PCR was performed for the gene expression using QPIG, QPPR, and ACTIN primers (Table S1).

### Immunoblot Analysis

Chloroplast membrane proteins were isolated from the leaves of around 2-week-old maize plants using kits (Cat no: BB-3175, BestBio, China). Protein samples were quantified with BCA protein assay. The protein samples were separated by 12% SDS-PAGE. After electrophoresis, the proteins were transferred onto a PVDF membrane (0.2 µm, Bio-Rad) using Bio-Rad Semi-Dry Transfer Cell. The blot was blocked with 5% milk in TBST for 1h at room temperature (RT) with agitation and then incubated in the primary antibody (from Agrisera, AS16 4065) at a dilution of 1: 500 overnight in +4°C. The antibody solution was decanted, and the blot was washed briefly with TBST at RT with agitation. The blot was incubated in secondary antibody (anti-rabbit IgG horseradish peroxidase conjugated, from Agrisera, AS09 602) diluted to 1:10 000 in 1% milk/TBST for 30min at RT with agitation. The blot was washed briefly in TBST at RT with agitation and developed for 2 min with ECL according to the manufacturer’s instructions (Cat no: SL1350, Coolaber, China). The signals were visualized by a GeneGnome chemiluminescence analyzer (Syngene).

## ASSESSION NUMBERS

Sequence data from this article can be found in the GenBank/NCBI databases under the SRA accession number: PRJNA588870.

## SUPPLEMENTARY DATA

**Supplemental Figure S1** Schematic representation of genotypes and kernel weights of recombinant families derived from F1H5.

**Supplemental Figure S2** qPCR analysis of Zm00001d048450 and Zm00001d048451 expression.

**Table S1** Primer sequences used in this study.

## ACKNOWLEDGEMENTS

This work was supported by the National Key Research and Development Program of China (2016YFD0100303), the National Natural Science Foundation of China (31525017, 31961133002), and NSF grant IOS-1546837 to DJ; Juan Huang was sponsored by China Scholarship Council to visit Cold Spring Harbor Laboratory and study in Prof. David Jackson’s lab from March 2018 to March 2019 (File No. 201706760027). We thank Dr. Yali Zhang from Shihezi University for suggestions on measurements of chlorophyll fluorescence. We thank Dr. Edgar Demesa Arevalo’s help in taking confocal images and for providing for P19 containing strain. We thank Dr. Hannes Claeys for providing pK7FWG2.0 containing strain. We thank Felix Fritschi for use of the chlorophyll fluorescence meter.

